# Spherical harmonics representation of the steady-state membrane potential shift induced by tDCS in realistic neuron models

**DOI:** 10.1101/2022.07.19.500653

**Authors:** Adrià Galan-Gadea, Ricardo Salvador, Fabrice Bartolomei, Fabrice Wendling, Giulio Ruffini

## Abstract

**Objective:** We provide a systematic framework for quantifying the effect of externally applied weak electric fields on realistic neuron compartment models as captured by physiologically relevant quantities such as the membrane potential or transmembrane current as a function of the orientation of the field.

**Approach:** We define a response function as the steady-state change of the membrane potential induced by a canonical external field of 1 V/m as a function of its orientation. We estimate the function values through simulations employing reconstructions of the rat somatosensory cortex from the Blue Brain Project. The response of different cell types is simulated using the NEURON simulation environment. We represent and analyze the angular response as an expansion in spherical harmonics.

**Main results:** We report membrane perturbation values comparable to those in the literature, extend them to different cell types, and provide their profiles as spherical harmonic coefficients. We show that at rest, responses are dominated by their dipole terms (*ℓ* = 1), in agreement with experimental findings and compartment theory. Indeed, we show analytically that for a passive cell, only the dipole term is nonzero. However, while minor, other terms are relevant for states different from resting. In particular, we show how *ℓ* = 0 and *ℓ* = 2 terms can modify the function to induce asymmetries in the response.

**Significance:** This work provides a practical framework for the representation of the effects of weak electric fields on different neuron types and their main regions—an important milestone for developing micro- and mesoscale models and optimizing brain stimulation solutions.

## 1. Introduction

Transcranial electrical stimulation (tES) is a non-invasive neuromodulatory technique based on accurate control of weak currents injected from multiple scalp electrodes and the resulting electric fields induced in the brain. tES, pioneered by Nitsche and Paulus [1], includes direct and alternating current variants known as tDCS and tACS. Low intensity, controlled currents (typically ~1 mA but ≤ 4 mA) are applied through scalp electrodes in repeated 20–60 min sessions. The subtle but persistent modulation of neuronal activity is believed to lead to plastic effects deriving from Hebbian mechanisms [2].

The current evolution of this and related techniques such as transcranial magnetic stimulation or deep brain stimulation is veering towards model-driven approaches and optimization principles based on a mechanistic understanding of pathophysiology, the interaction of the electric field with neuron populations and the resulting dynamical, and then plastic, effects [3, 4]. In this work, we aim to quantify in a systematic way the response of different cell types and cell parts to be used in future approaches to model-driven stimulation.

The concurrent effects of tES currents on the brain are thought to be mediated by the generated electric field’s action on cortical neurons [5, 6]. These electric fields are weak (~1 V/m) [7], relatively large-scale (with spatial correlation scales of the order of a few cm), and induce small changes in the membrane potential of cells (sub-mV) [8, 9]. Polarization happens in a compartment-specific manner [10, 11, 12], hyperpolarizing compartments close to the virtual anode and depolarizing those close to the virtual cathode. Furthermore, it is expected from cable theory that changes in the electric field along neuron fibers cause local polarizations as well. Then, the effect in a cell under the same stimulation can have very different responses locally. The effects are significantly larger where fibers bend and mostly at terminations [13]. Thus, the overall morphology is key to explaining effects observed locally. To take all effects into account, we use multi-compartment models that allow us to locally couple the field to realistic morphology models and make predictions of the induced perturbations [14].

Pyramidal cells have been of particular interest in the field for two reasons. First, they are elongated, which leads to the generation of a larger membrane perturbation when the field and neuron are aligned [9]. Secondly, they are spatially organized. Because of this and the spatial homogeneity of the electric fields, the effects on such cells are spatially coherent and thought to lead to stronger, network-enhanced effects [15, 16]. In contrast, effects on interneurons are usually disregarded due to their small size and non-elongated branching profiles [9].

In this paper, we focus on characterizing the effects of the weak electric field on the stable equilibrium of single neuron models using expansions in terms of spherical harmonics. With this approach, we aim to provide (i) a systematized method for describing the angular dependence of the response through simulations, (ii) a framework that allows for the averaging of the angular response of many compartments to describe cell parts and types, and (iii) a representation in meaningful terms that represent the dipolar behavior on its own but accounts for other angular dependence profiles if necessary. We validate the common dipolar approach and that it is accurate for small perturbations induced at neurons at rest (far from the threshold) and show that it is exact for the passive membrane. Nevertheless, we observe that corrections to pure dipolar responses may need to be applied closer to the threshold in realistic neurons.

## 2. Methods

### 2.1. Formalism for steady-state changes induced by an electric field

We will be studying the response of a neuron to a constant, spatially uniform, weak electric field as typically generated by non-invasive brain stimulation methods such as tDCS — at least away from tissue boundaries, since these can introduce strong local changes in the field distribution due to the change in electrical properties between tissues. The field is characterized by a magnitude *E* ≡ ∥**E**∥ and an orientation that can be expressed with two spherical angles, the polar angle *θ* and the azimuth *φ*. Thus, in spherical coordinates, the electric field is given by

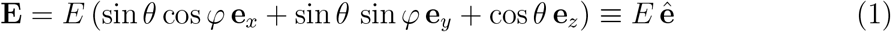

Let us consider a variable *v* that characterizes a physiologically relevant magnitude affected by an external electric field. Our study, focuses on the membrane potential, but the formalism can be extended to other variables, e.g., transmembrane currents. If the variable v has a stable equilibrium, i.e., resting potential in our case, we expect that the applied electric field will shift that equilibrium to a new value *v′*. We aim to quantify that induced change *δv* of the steady-state solution for different realistic models of cell types. In our case, this shift is commonly referred to as polarization.

Let us assume that two independent terms can express the effect of the magnitude and orientation. On the one hand, the polarization *δv* scales linearly with E, which has been observed experimentally for small fields [9]. Regarding the dependency of the shift to the orientation of the field, we propose a function defined on a sphere Φ(*θ, φ*), i.e., that has a value for any direction in space. We refer to Φ as the *response function,* which is a function that outputs the shift *δv* under an electric field of 1 V/m in a given direction. Under these assumptions §, *δv* can be expressed as shown in Equation 2.

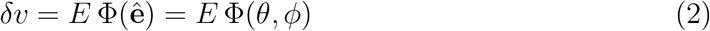

### 2.2. Expansion in spherical harmonics

Spherical harmonics form a complete and orthonormal set of functions *Y_ℓm_* defined on the surface of the sphere (parametrized by two spherical angles, *θ* and *φ*), where *ℓ* and *m* refer to the degree and the order of the harmonic, respectively. Squared-integrable functions defined on the sphere can be expressed as a sum of spherical harmonic terms [17, 18], much as functions on the plane can be expressed as a Fourier series. In fact, both spherical harmonics and complex exponential are eigenfunctions of equations associated with the Laplace partial differential equation in different manifolds (the plane and the sphere). As the response function Φ(*θ, φ*) is a function on the sphere, it can be expanded in terms of spherical harmonics, meaning that it can be expressed as in Equation 3 and fully characterized by a set of coefficients *f_ℓm_*,

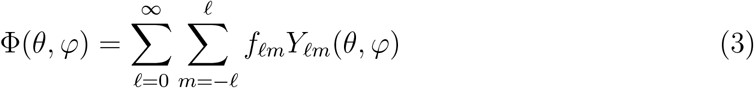

Spherical harmonics are characterized by a degree and order pair (*ℓ, m*); see Appendix A for a definition. They are tightly linked to the multipole expansion, so one can think of the *ℓ* =0 term as a *monopole* behavior, a term that does not depend on orientation. The *ℓ* =1 terms represent a dipolar behavior, *ℓ* = 2 a quadrupole, and so on. Finally, under a parity transformation, where a point with coordinates {*θ, ϕ*} changes to {*π* – *θ, π* + *ϕ*} (corresponding to a reversal of the orientation of the electric field in our case),

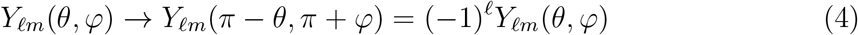

This means that if we expect the membrane response to change sign under electric field reversal (e.g., if it is linear in **E**), only odd *ℓ* terms should contribute to the expansion.

#### The l-2 norm of the coefficients is an influenceability measure

The *l*-2 norm of the coefficients quantifies the maximum of the response function regardless of the orientation. The actual values are subject to the specific normalization used. We name this measure the influenceability *ξ* of the response function,

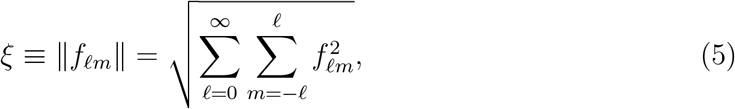

and serves as a metric to compare how affected in magnitude by an electric field each model is. In other words, for a set of response functions *f_ℓm_,* we can compare how susceptible they are to being influenced given the same electric field.

#### The λE linear model is contained in the ℓ =1 terms of the expansion

The *f*_1*m*_ terms correspond to the *λ* (dipole) in the *λ*E model, where the induced polarization is represented as the dot product of the electric field vector **E** and a vector *λ* of magnitude the length constant of the membrane and in the orthodromic direction of the cell [5]. We can rewrite the *λ*E model as

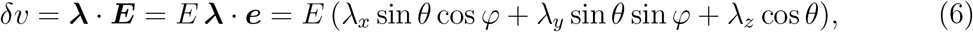

which is seen to correspond to the *ℓ* =1 terms in the expansion, with *λ_z_* is represented by *f*_10_, *λ_x_* by *f*_11_ and *λ_y_* by *f*_1-1_. Thus, the *λ*E model predicts symmetric polarizations. In particular, inverting the direction of **E** inverts the polarization pattern. This translates to polarizations equal in magnitude but opposite in sign in anodal and cathodal conditions, as shown in Figure 1.

**Figure 1.**
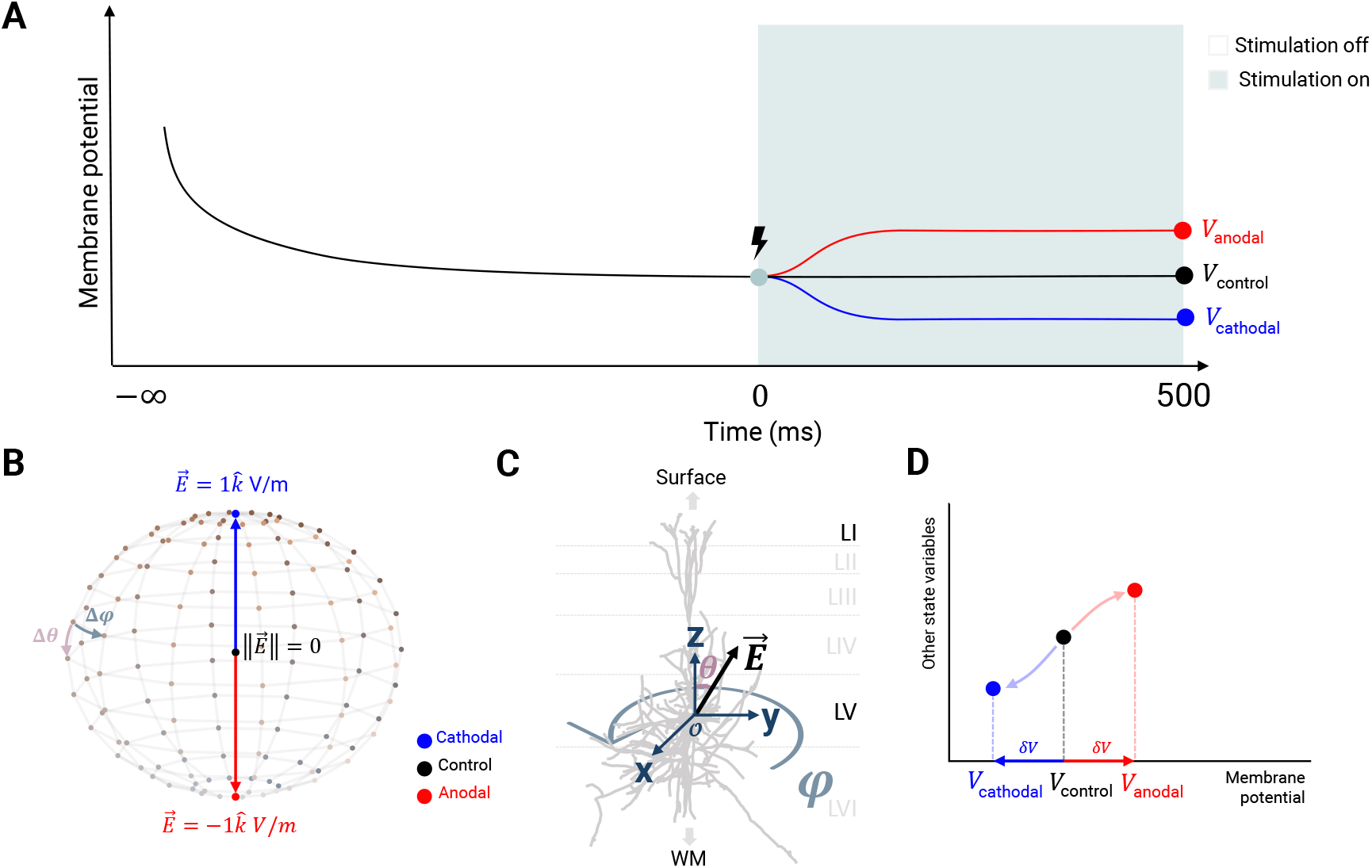
Experimental design for estimation of the response function. **A. Temporal profile** of the simulation of different experimental conditions. The first part of the experiment computes the resting condition of the model and is common for all experimental conditions. The latter consists in applying a constant external electric field and is repeated of all the desired orientations of the electric field. **B. Experimental conditions.** Anodal (red), cathodal (blue) and control (black) conditions in the context of the DH1 Sampling of the sphere. The grid represents the *N×N* sampling of the sphere with *N* = 12, with points separated by Δ*θ*=*π*/13 and Δ*φ* = 2*π*/13 [23]. **C. Coordinate system** definition with respect to the cell. The vertical direction represents the component normal to the cortical surface and corresponds to the *z* Cartesian coordinate. The spherical coordinates represent with *θ* the inclination and with *φ* the azimuth. The radius is the magnitude of the field *E*. For all cell models, the origin of coordinates *O* is set at the center of the soma. **D. Stimulation as an attractor shift.** A constant applied field changes the steady-state condition. In the soma of a prototypical pyramidal cell, the state is shifted towards a depolarized membrane potential.

### 2.3. Cell models

We use the multi-compartment conductance-based models from the cortical microcircuit reconstruction by the Blue Brain Project [19]. There, model morphologies are based on reconstructions from real acquisitions from juvenile rats. This work, considers all the morphology types in the database except the layer I interneurons. That includes nine types of interneurons: bipolar, bitufted, chandelier, double bouquet, Martinotti, neurogliaform, and basket cells (including large, small, and nest). These nine types make 36 morphology types, one of each for each layer (II/III, IV, V, and VI). Regarding excitatory cells, there are 13 morphology types (12 types of pyramidal cells and spiny stellate cells). All morphologies are made of connected cables, as required by the NEURON simulation environment (v8.0.0). Each cable is discretized in evenly spaced compartments. All cables were assigned a minimum of one compartment and, two compartments were added every 40 μm of length — as in the original work. The biophysical membrane models contain 13 different known Hodgkin-Huxley type ion channel models distributed using different criteria along the morphology. The Blue Brain Project team defined 11 ion channel profiles or *electrical types* exhibiting different spiking features. The options unfold into ten inhibitory and one excitatory electric profile, where the latter is the one representing all excitatory cell models of the dataset (continuous adapting, cAD). Regarding inhibitory cells, they included bursting (b), continuous (c), and delayed (d) models, later subdivided into accommodating (AC), non-accommodating (NAC), stuttering (STUT), and irregular (IR). Irregular spiking models contain stochastic potassium channels that introduce stochasticity into their behavior. For this study, we suppressed these because of incompatibility with our framework. Along the same lines, we discarded electric types generating spontaneous spiking at rest: cNAC, cSTUT, dSTUT, and dNAC. Given that the same morphology types were found to show different electrical types, they created models for each morphoelectrical combination. Then, the model dataset has five statistical clones of each morphoelectrical type.

#### 2.3.1. Uniform electric field coupling

The coupling of the external electric field to the neuron models is implemented via the extracellular potential in NEURON 8.0.0. We use the extracellular mechanism to access the `extracellular` potential in each compartment. To attach the value induced by the external electric field, we tweak the xtra mechanism for extracellular recording and stimulation based on the approach by McIntyre and Grill [20]. Recall that the extracellular potential is related to the external uniform field by **E** = –**▽***V*_ext_, with

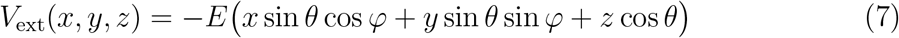

Here (*x, y, z*) refers to the spatial coordinates in 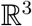 of the center of a compartment of the neuron model.

### 2.4. Response function estimation

Let 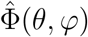 be the discretized estimation of the response function. 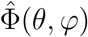 can be defined as a collection of changes *δv* observed for different orientations, where the particular orientation set is determined by the sampling method and the chosen resolution. We obtain 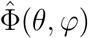 performing simulations applying electric fields of 1 V/m of magnitude in different orientations to a resting neuron model using the NEURON simulation environment (v8.0.0) [21]. All experiments are performed at temperatures of *T* = 36.9°C to mimic the average physiological temperatures of living human brains [22].

#### 2.4.1. Temporal design of the simulation experiment

The simulation pipeline to estimate the steady-state response of any neuron model that has a stable equilibrium under an electric field consists of two phases: (1) computing the resting state and (2) applying an electric field to estimate the change in the induced change in the steadystate condition. The resting state is defined as the steady-state solution of the system when no external stimulation is present. It is assessed by letting the system converge until it experiences no further change in its state variables. Computationally this requirement is met when the membrane potential varies less than 1 nV anywhere in the cell without further external inputs. The second phase assesses the steady-state solution’s displacement under an external perturbation; see Figure 1.D. The system is initialized in the pre-computed resting state. Then, the shift in the steady-state solution is retrieved as the change experienced after 500 ms of 1 V/m uniform electric field application in a given condition. We chose 500 ms as a relatively large fixed simulation time at which we can assume the system has reached a stable solution. The induced change, i.e., *δv*, is computed with respect to the control condition with no field applied. All simulations are integrated using the fully implicit backward Euler method with an integration time step of 25 μs.

#### 2.4.2. Experimental conditions

To characterize the response function, we assess the induced changes for different orientations. We use the Type I Driscoll-Healy sampling of the sphere (DH1), using *N* =12 to reconstruct harmonics until a maximum degree of *ℓ*_max_ = 5. The sampling theorem gives the relationship among both: *ℓ*_max_ = *N*/2–1 = 5. The DH1 sampling consists of sampling both *θ* and *φ* in *N* points each, building an *N×N* grid covering the sphere [23]. Because the full span of both angular dimensions is used, the polar resolution (Δ*θ*) is half the azimuthal one (Δ*φ*). For *N* = 12, the angular distance among consecutive sampled points is Δ*φ* = 2*π*/13 and Δ*θ* = *π*/13 rad. The discretization is shown in Figure 1.B.

#### 2.4.3. Spherical harmonics-based representation

Following the expansion introduced in Equation 2, once 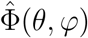 has been estimated for each compartment, it is represented by a set of spherical harmonics coefficients. The calculation is performed with the pyshtools Python package, using a 4*π*-normalization and the Condon-Shortley phase factor convention [24]. The coefficients *f_ℓm_* fully represent the response function Φ(*θ, φ*), which can be assessed by integrating the response function over the sphere. Using the 4*π* normalization, it reads as a double integral over the sphere for a continuous Φ(*θ, φ*).

In our case, it translates to a discrete sum, as in Equation 8.

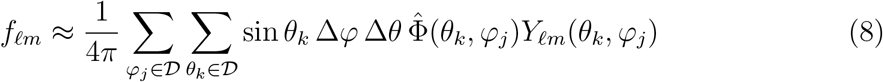

where 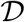 is the domain of sampled points. Modern implementations of these expansions use fast Fourier transforms over latitude bands, among other methods, to improve computational efficiency [25, 24].

### 2.5. Averaging response functions

A response function can be derived for each compartment of a cell model through the methodology already described above, v. Figure 1. However, in the context of model-driven stimulation, the interest lies in simplifications of the complex morphology. Due to linearity, *lumped* response functions can be obtained by averaging the coefficients. We exploit this to create two models: response functions of (i) cell parts and (ii) cell types. The multi-compartment models here have hundreds of compartments for each cell along their morphology. Each compartment can be tagged to belong to a specific region or part of the cell. We use the model’s original classification of axon, soma, and dendrites, with a further distinction of apical and basal dendrites in the case of pyramidal cells. In the case of apical dendrites, we make a further distinction between those found in LI with respect to the rest. To build models of cell types, we use the different statistical clones available in the database belonging to the same cell type. We averaged the models of the different cell parts of the clones of the same type and, this way built a canonical model for that type.

### 2.6. State-dependence

Response functions are expected to vary depending on the morphology and biophysical models of the cells. The ion channels models are functions of the membrane potential and hence depend on the state of the cell. Thus, we expect that response functions are also state-dependent, and we set out to investigate how it may vary for the soma of the canonical LV thick-tufted pyramidal and the size of these effects. Given our steady-state analysis framework, we reconstruct a response function for different membrane potential values using a current clamp at the same study compartment, i.e., the soma. We apply injected currents from −0.2 nA until the system loses its stable equilibrium and stops converging — i.e., the neuron model starts firing. The membrane potential also affects the time constants of ion channel models, and they are expected to increase when moving closer to spiking regimes. For this reason, we provide the system for longer times in experiment phase 2 (10 s) to assess the steady-state change. Currents are injected prior to the application of the electric field and kept during the whole simulation. In other words, we change the location of the stable equilibrium through an external current and explore the perturbations induced by a 1 V/m electric field in that new vicinity.

## 3. Results

### 3.1. The response function at rest is dipolar

Consistent with the analysis of the passive cell (see Appendix B), the response function is dipolar under weak fields at rest. In other words, only *ℓ* =1 terms are nonzero. The example of the response function of the soma of a pyramidal cell model is illustrated in Figure 2, where 2.C shows the different terms with only the term *ℓ* = 1, *m* = 0 being nonzero. All coefficients of all cell models analyzed are shown in Figure ??. During stimulation, each cell compartment reaches a new steady-state solution. The maximum observed somatic polarization corresponds to fields parallel to the somatodendritic axis at 0.14 mV per 1 V/m in layer V early-branching thick-tufted pyramidal cells. These values are in line with the literature as with results reported in CA1 pyramidal cells [26] and comparable to the typically assumed value of 0.2 mV per V/m of applied electric field [9].

**Figure 2.**
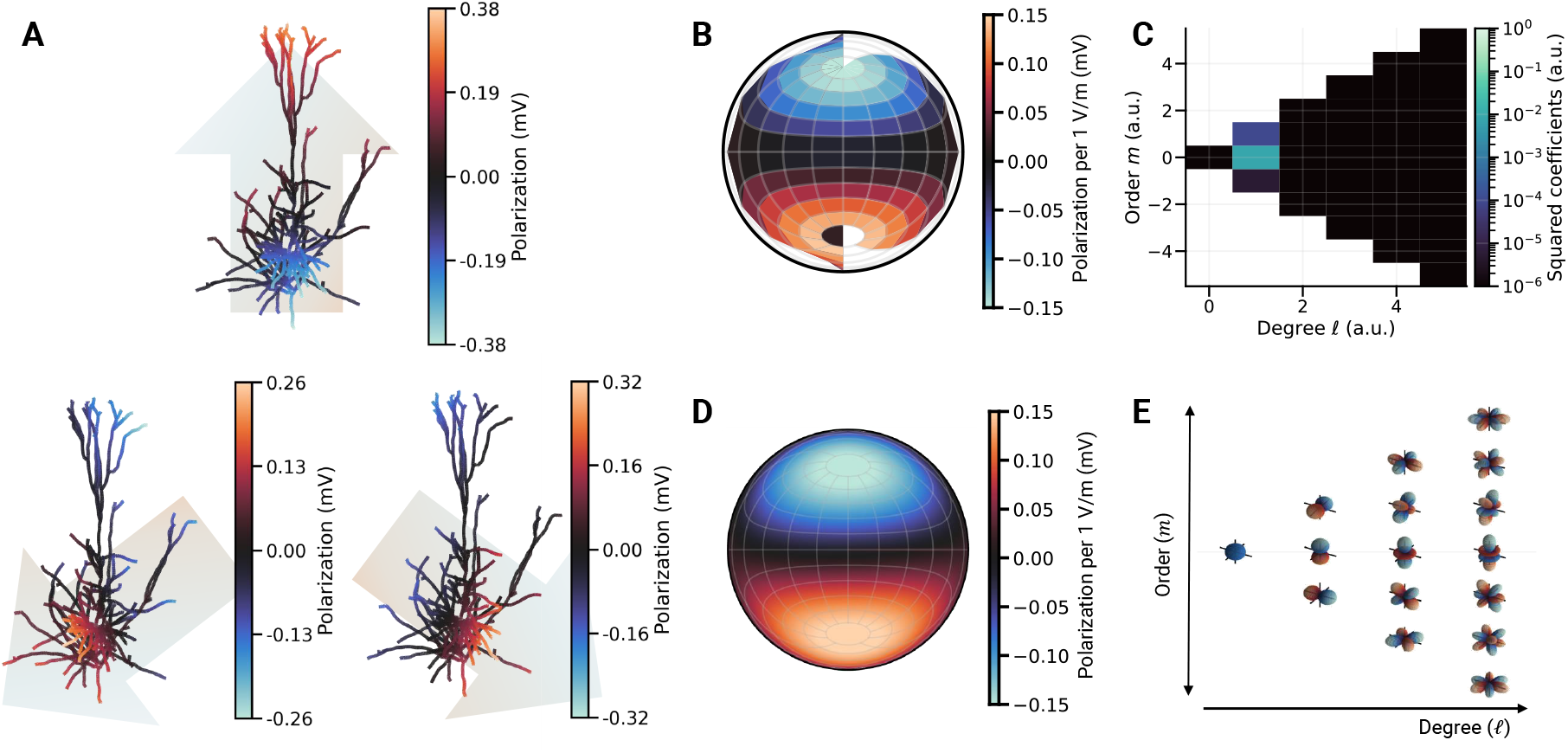
Polarization dependence on the electric field orientation. **A. Polarization of the membrane under a uniform electric field of 1 V/m.** The direction of the external electric field has a direct impact on how the charges are redistributed along the cell morphology. In this example, the effects on a layer V pyramidal cell model are displayed for an applied external electric field of 1 V/m in the direction of the background arrows displayed below each cell. **B. Somatic response function estimate.** Estimated polarization of the soma under a 1 V/m electric field for each orientation of the DH1 sampling as described in Figure 1.B. The 2D projection is used with the Lambert method, where the azimuthal angle matches the horizontal axis. **C. Spherical harmonics spectrum.** Representation of each spherical harmonic coefficient squared. Only the *ℓ* = 1 terms are present, i.e., the response is dipolar. The description in spherical harmonics terms displays a strong predominance of the (*ℓ* = 1, *m* = 0) indicating an alignment with the z-axis (see Figure 1.C) slightly tilted towards the x-axis as determined by the presence of a small (*ℓ* = 1, *m* = 1) term. **D. Reconstructed response function.** Continuous version of the response function represented by the coefficients. It represents all the information contained in the sampled estimate shown in B. **E. Spherical harmonics.** Polar representation of all spherical harmonics up to *ℓ* = 3 in the same layout as in C for an intuitive interpretation of each of the coefficients. The radius codes the magnitude of the function for each orientation and the color codes the sign, warm and cold being positive and negative respectively.

### 3.2. Somatic polarizations in excitatory cells

Consistent with previous findings, the most affected cells are pyramidals of subgranular layers (V and VI), with an increased effect on those with an apical tuft. The most considerable effect is observed in the early-branching thick tufted population due to the asymmetry of the processes distribution with respect to the soma. Different responses also accompany the variety of layer VI pyramidal cells. Somatic polarization in the LVI tufted cell models is at 0.10 mV, i.e., 58% larger than in the untufted, at 0.07 mV. Somata of inverted pyramidals show an effect of size comparable to that of tufted cells, i.e., 0.10 mV but with reversed polarity. In supragranular layers, effects on pyramidal cells were minimal and irrelevant on spiny stellate and star pyramidals. All somatic polarizations for the different averaged pyramidal cell models are shown in Figure 3 and their maximums together with their respective ratios are compiled in Table 1. The table also features the 95^th^ percentile of the maximum polarizations observed within cell before averaging as a more agnostic statistic of the polarizations reached throughout the morphology.

**Figure 3.**
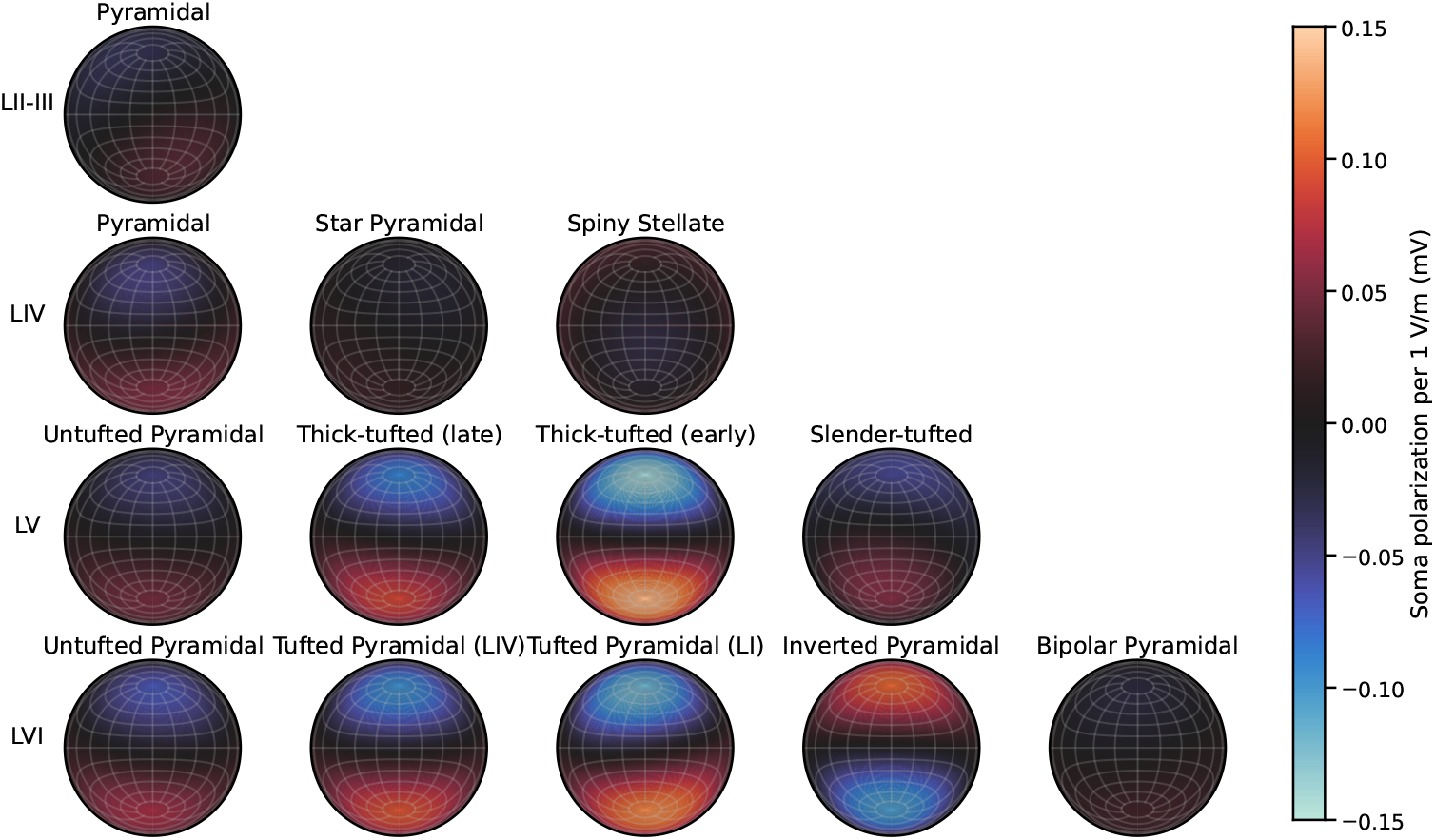
Polarization of soma of pyramidal cells under 1 V/m electric field. Each row represents a neocortical layer and each column a pyramidal cell type. Each plot displays the modeled change of the membrane potential (mV) in the soma due to an external electric field of 1 V/m as a function of the relative orientation of the electric field. Orientation dependence is represented using a Lambert projection. The vertical axis codes the polar angle *θ* which ranges from *θ* = 0 in the top (indicating a vertically aligned electric field pointing up) to *θ* = *π* (down) in the bottom. The horizontal axis is the azimuthal angle *φ*, where the left end is *φ* = 0 and the right one is *φ* = 2*π*. “*Early*” and “*late*” in layer V (LV) thick-tufted pyramidal cells refer to when the apical dendrites branching of the tuft begins. In LVI tufted pyramidal cells, the layer in parenthesis is the layer in which the dendritic tuft terminates.

**Table 1.** Maximum observed somatic (S) polarizations *δV* and transmembrane currents *δI_m_* rounded to the μV and fA, respectively, and their respective ratio referenced to the largest. Additionally, the 95^th^ percentile (P95) of the maximum polarization observed compartments is reported too. The cell type with the largest effect is the LV thick-tufted cell with an early branching of the apical tuft (bold).

| Cortical layer | Cell Morphology type | $\delta V$ (S) | $\delta V$ (P95) | $\delta I_m$ (S) | Ratio | Ratio |
| --- | --- | --- | --- | --- | --- | --- |
| | | ($\mu V$) | ($\mu V$) | (fA) | $\delta V$ (S) | $\delta I_m$ (S) |
| LII/III | Pyramidal | 36 | 251 | 26 | 0.26 | 0.11 |
| LIV | Pyramidal | 52 | 232 | 27 | 0.38 | 0.11 |
| LIV | Star pyramidal | 19 | 200 | 5 | 0.14 | 0.02 |
| LIV | Spiny stellate | 25 | 204 | 14 | 0.18 | 0.06 |
| LV | Untufted pyramidal | 49 | 189 | 28 | 0.35 | 0.12 |
| LV | Thick-tufted pyramidal (late) | 89 | 238 | 122 | 0.64 | 0.52 |
| **LV** | **Thick-tufted pyramidal (early)** | **138** | **264** | **233** | **1.00** | **1.00** |
| LV | Slender-tufted pyramidal | 56 | 211 | 65 | 0.41 | 0.28 |
| LVI | Untufted pyramidal | 68 | 199 | 21 | 0.49 | 0.09 |
| LVI | Tufted pyramidal (LI) | 96 | 188 | 36 | 0.70 | 0.16 |
| LVI | Tufted pyramidal (LIV) | 119 | 199 | 30 | 0.87 | 0.13 |
| LVI | Inverted pyramidal | 104 | 208 | 27 | 0.76 | 0.11 |
| LVI | Bipolar pyramidal | 25 | 200 | 9 | 0.18 | 0.04 |

### 3.3. Somatic polarizations in interneurons

Due to its central position with respect to other cell processes, effects in the soma of interneurons are rather small compared to pyramidal cells. Restricting the electrical type to cAC models, we observe a large variety of responses across morphologies and layers—see Figure 4. In bipolar, bitufted, chandelier, and double bouquet cells, we observe a reversal of the polarity of the effect at the soma between supragranular and infragranular layers. Martinotti cell models show the lowest variability between layers. Basket cells are the group that displays the smallest response at the level of the soma consistently in their different types (large, nest, and small) and layers. On the other hand, the cell type model displaying the largest somatic effect is bipolar cells. In this particular cell morphology, the size of the effect strongly depends on its electrical type. However, we do not observe such dependence in the other types of interneurons. The responses across layers and for different electrical type models of bipolar cells are shown in Figure 5, where it is observed how the effect induced in the soma is reversed in lower layers compared to upper ones and how the electrical type models influence the effect size.

**Figure 4.**
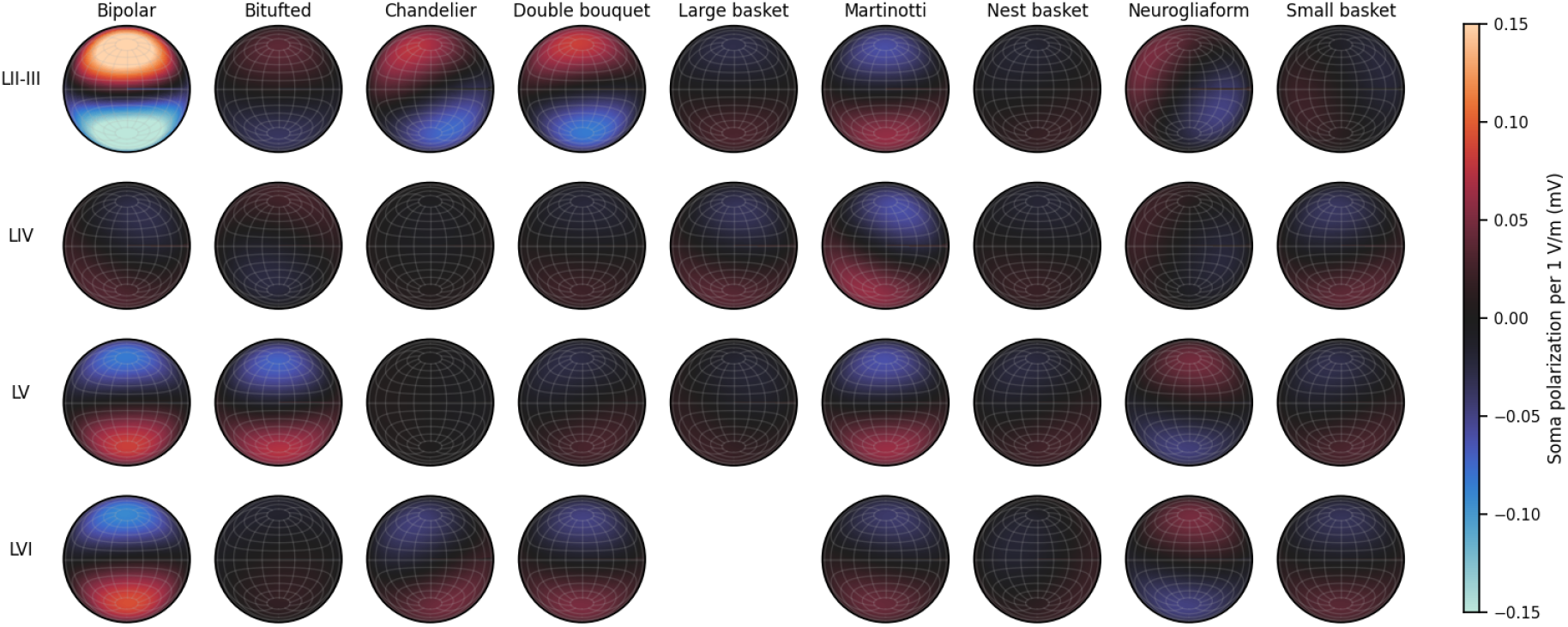
Polarization of soma of interneurons under 1 V/m electric field. Simulations of the average membrane potential change (mV) in the soma in continuous accomodating (cAC) models of interneurons. Rows and columns represent cortical layers and morphology types respectively. The color range is set to match that of Figure 3 to ease comparison with pyramidal cells. Bipolar cell models of layers II-III appear saturated because their polarization values exceed the 0.15 mV. Large basket cell models for LVI are not available in the original set of models.

**Figure 5.**
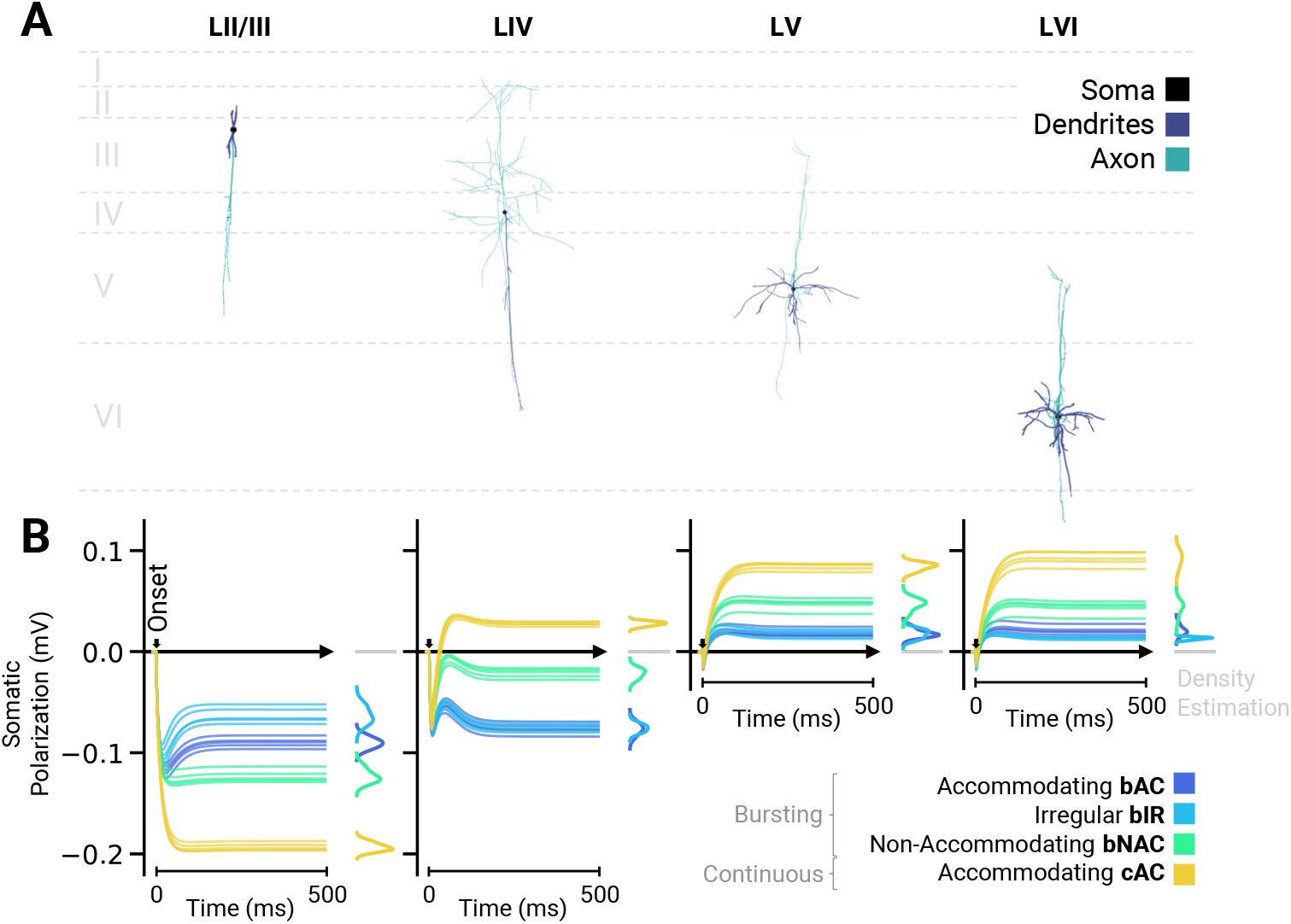
Effect differs across electrical types of bipolar cells. **A.** Morphology examples of bipolar cells of different layers. **B.** Somatic responses of the different cell models under anodal stimulation of 1 V/m. Each plot provides the polarization curves as a function of time of the models of bipolar cells of a given layer. From left to right, they correspond to LII/III, LIV, LV and LVI models following the outline of A. Each line is the trace of the soma of a different cell model, where the color codes the electrical type of the model. There are five models per electrical type. The stimulation onset is set at time equals zero and is marked as a small arrow over the horizontal axis. The total stimulation duration is of 500 ms. At the right of each plot, the distributions — probability density estimations — of steady-state polarizations reached at the end of the 500ms for the different electrical types are shown. Colors are set to match to those of the lines, see legend at the bottom right part of the plot.

### 3.4. Effects on different cell parts can be represented with different response functions

As expected from the pre-existing literature on the field, different parts of pyramidal cells respond differently to the presence of the field. Analyzing the effect locally, i.e., compartment-wise, bends and fiber terminations increase the effect of the field on membrane polarizations. The overall size of the this effect is captured by the influenceability metric as defined in Equation 5. In a pyramidal cell, we observe how the most susceptible areas are the tuft of the apical dendrites and terminations at the axon and the basal dendrites, see Figure 6.

**Figure 6.**
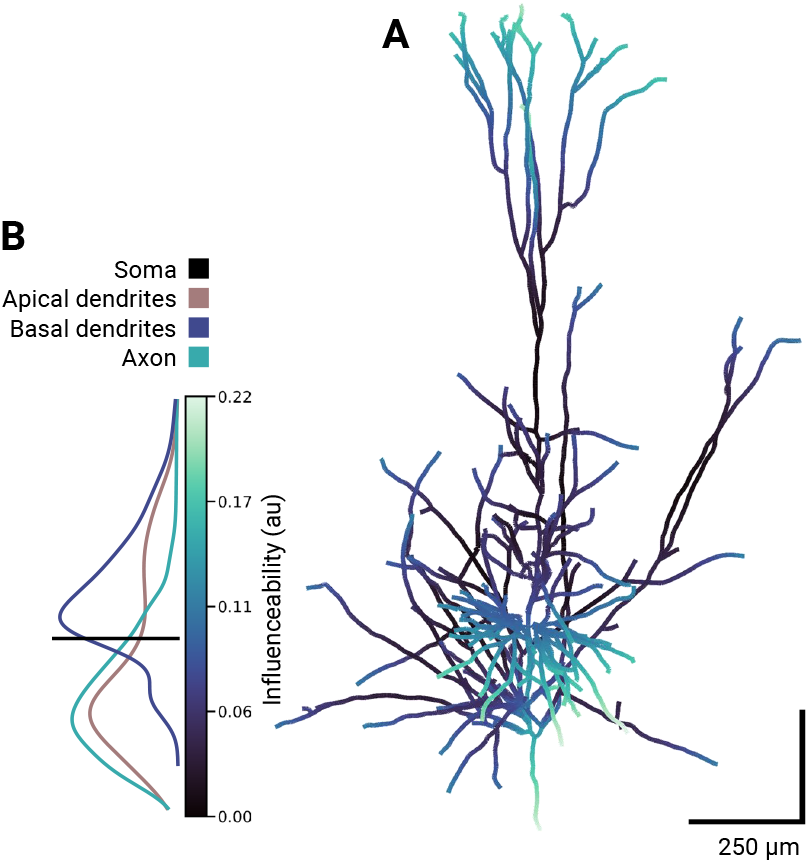
Influenceability values of a pyramidal cell. **A.** Variation of the influenceability measure throughout the morphology of an early-branching LV thick-tufted pyramidal cell. Scale: 250 μm. **B.** Distributions across compartments associated with different cell parts. The distribution of the apical dendrites displays bimodality, distinguishing the difference in values along the apical trunk from the values observed in the apical tuft.

On the other hand, we provide functions for the major regions of the cell. The estimated response functions for the different cell parts, obtained through averaging the responses of the individual compartments belonging to each region, display the classical response of the apical dendrites with a reversed polarity compared to the one observed in the soma and the basal dendrites, e.g., hyperpolarized under anodal stimulation, see Figure 7. The largest response among the explored regions is found in the apical dendrites contained in layer I. In the example of the figure, the polarization in the direction where maximum effect reaches 0.24 mV per V/m in layer I apical dendrites, whereas it is only 0.03 mV at the trunk, defined as the compartments from the apical tree found from layer II down. When averaging the whole of the apical dendritic tree, the response function is reduced to a maximum of 0.07 mV (per V/m). The soma and the basal dendrites display a similar pattern, with a maximum polarization of 0.16 in the soma and 0.18 mV in the basal dendrites. In the axon, including the descendant and the collaterals, the average is smaller at 0.04 mV per V/m.

**Figure 7.**
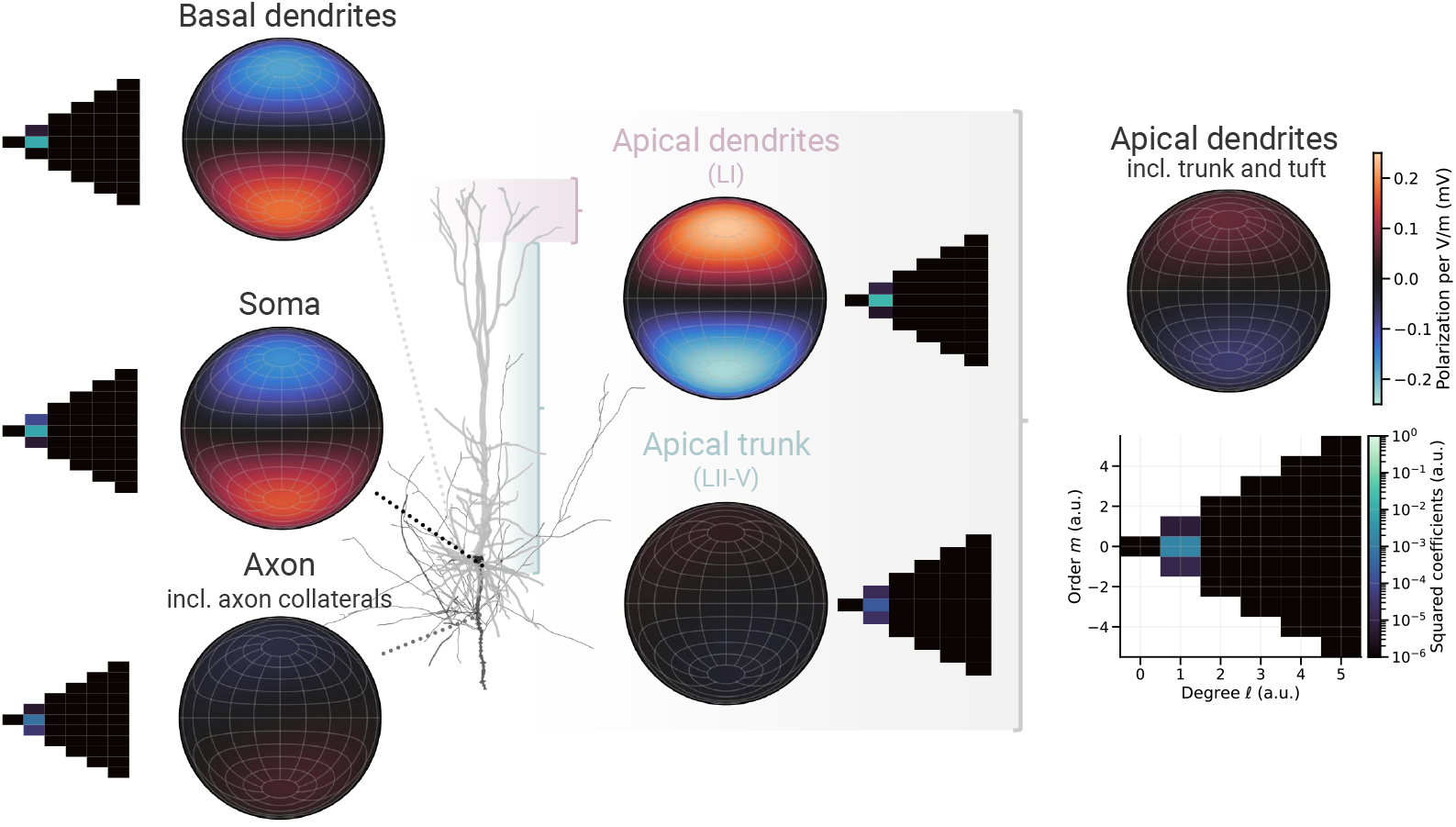
Models of different sections of a cell. Average response functions of different cell parts of a layer V (LV) early-branching thick-tufted pyramidal cell model and their 2D spectrums. In the morphology, the different cell parts considered are shown in different shades of gray. Models of apical dendrites are split in compartments belonging to L1, to the rest of layers and to the whole of the apical arborization. Coefficients are reported in Table ??.

### 3.5. The direction and magnitude of the effect depend on the cell state

We observe that the response function changes by injecting hyperpolarizing and depolarizing currents in the soma of the layer V pyramidal cell model. Recall that the injection starts before applying the electric field, and the system is already in its corresponding steady state, given the injected current. It keeps displaying a dipolar behavior for most values, purely captured by the *ℓ* =1 coefficients. Moreover, the most relevant term in all the states is the dipole in the direction parallel to the normal to the cortical surface, i.e., aligned with the z-axis (*ℓ* = *1,m* = 0) as in Figure 1.C. Under a hyperpolarizing current, we observe a slight increase of the effect saturating at 0.17 mV per V/m. Under depolarizing currents, we observe two different types of behavior.

For currents up to 0.25 nA (corresponding to a membrane potential of ~-62.50 mV, 10 mV above resting), we observe a decrease in the magnitude of the effect reaching a minimum of 0.12 mV per V/m in the direction of maximum effect. We observe a different behavior for currents above 0.25 nA until the system enters a spiking regime for currents above 0.34 nA, corresponding to potentials over −58.13 mV. In this window, we observe a rapidly increasing effect and the emergence of a non-dipolar response. The response becomes asymmetric, i.e., the magnitude of the effect of anodal and cathodal stimulations of equal field magnitude is not the same. Still, fields perpendicular to the axis of the *dipole* have negligible effects. In spherical harmonics terms, we observe it as a pair of *ℓ* = 0 and *ℓ* =2 terms that increases with the size of the asymmetry, being largest at the verge of the bifurcation. Right before the bifurcation, we report an effect of anodal stimulation of 0.35 mV, whereas cathodal stimulation shifts the membrane potential by −0.26 mV. The inducible changes thus span a range of 0.61 mV with an asymmetry of 0.09 mV.

## 4. Discussion

One of this study’s primary goals is to develop a simple and intuitive framework that quantifies the effects of weak fields that can be used to build large-scale models. The values shown here for the juvenile rat reconstructions serve as a proof of concept that can be extrapolated to cell models of other species or even the same ones adapted to match dimensions and other physiological features to that of adult rats or humans [14]. The approach applies to any multi-compartment model with a precise enough spatial resolution in its morphology.

### Modeled effects from a dynamical systems perspective

Equation 2 quantifies the displacement of the stable equilibrium of a neuron model under a weak electric field. As previously discussed, the model requires this type of solution in the analyzed regime. However, neurons are excitable because they operate close to bifurcations that change the dynamical landscape, generally to give rise to periodic solutions, e.g., sustained firing. For instance, integrator models have a stable node (resting) and a saddle node. Short excitatory inputs push the system closer to the saddle-node from the stable node, and if the system is pushed over the stable manifold of the saddle (threshold), it elicits an action potential on its excursion back to the stable point. A constant electric field works by persistently displacing the stable solution, and that is what the framework proposed quantifies: the shift of the stable node. In the example of an integrator model, this shift can be towards (excitatory) or away from (inhibitory) the spiking threshold. However, if the system is in an oscillatory regime, our formalism enters a problem of definition and cannot be strictly used anymore. A formalism in similar lines should be developed to describe changes in oscillatory solutions to model effects of rhythmic fields of similar sizes — like those generated by tACS or endogenous ephaptic interactions, which allow different modulations by entraining synchronizations [27].

### The leading terms in the spherical harmonic expansion are the dipolar ones

The expansion in spherical harmonics of the angular dependence of the effect provides a general framework containing different elements of interest in its different terms. Relevant to stimulation models based on head models, in which one cannot know the azimuthal orientation of a cell with respect to the electric field induced, it becomes convenient that the expansion already collects in the *m* = 0 terms the behavior that does not depend on the azimuth. In fact, by definition, the *m* = 0 terms alone describe the *average* behavior across all azimuths. Another relevant feature is that the method quantifies the dipolar part of the response in its *ℓ* =1 terms, which we find to be the only present for most cases, in agreement with theoretical arguments (Appendix B). This suggests that after the quantification, the steady-state shift can be represented by a dot product as proposed in the literature under the *λ*E model framework [5]. Furthermore, in realistic settings where azimuthal information of single neurons is unavailable and for cells near rest, the only relevant term is thus the (1,0).

However, we have observed how dipoles are not the only type of response, and we observe asymmetries among anodal and cathodal responses while keeping a neutral response to perpendicular fields in cells closer to the threshold. This behavior needs other terms that modify the dipolar ones accordingly. For dipoles aligned with the z-axis (1,0), we observed the emergence of a pair of equal values of (0,0) and (2,0). This pair shifts the values of the 1-degree terms at the poles of the dipole. Equivalent expressions involving other terms can modify dipoles oriented in other directions.

### State-dependence

In the neighborhood of the resting condition, we observed an increase in the effect size if currents were hyperpolarizing and a decrease if depolarizing, as seen in Figure 8.A. This effect aligns with the fact that the size of the polarization induced by the external electric field depends on the membrane resistance. If the resistance increases, the effect’s magnitude should also increase [28]. Nevertheless, there are opposite views in this regard since it has also been pointed out that active networks — with lower membrane resistance on average, as supposedly ion channels are opening more to fire — are preferentially modulated in tDCS [29]. The increased excitability we observe close to the bifurcation seems to be more cohesive with the latter view, which reads as if the system is placed very close to the critical point by receiving more input, the effect appears larger. On the other hand, the window of increased excitability right before the bifurcation is also consistent with the typical effect of tDCS affecting spike times [30]. A large effect in the vicinity of the emergence of spikes can underlie a modulation of the moment at which a spike happens. This phenomena has been postulated as a possible mechanism of action of tES via spike time-dependent plasticity [31, 32]. The emergence of other terms, i.e., asymmetries and increased excitability towards the spiking threshold, cannot be explained just by a passive-like membrane argument and requires a different interpretation.

**Figure 8.**
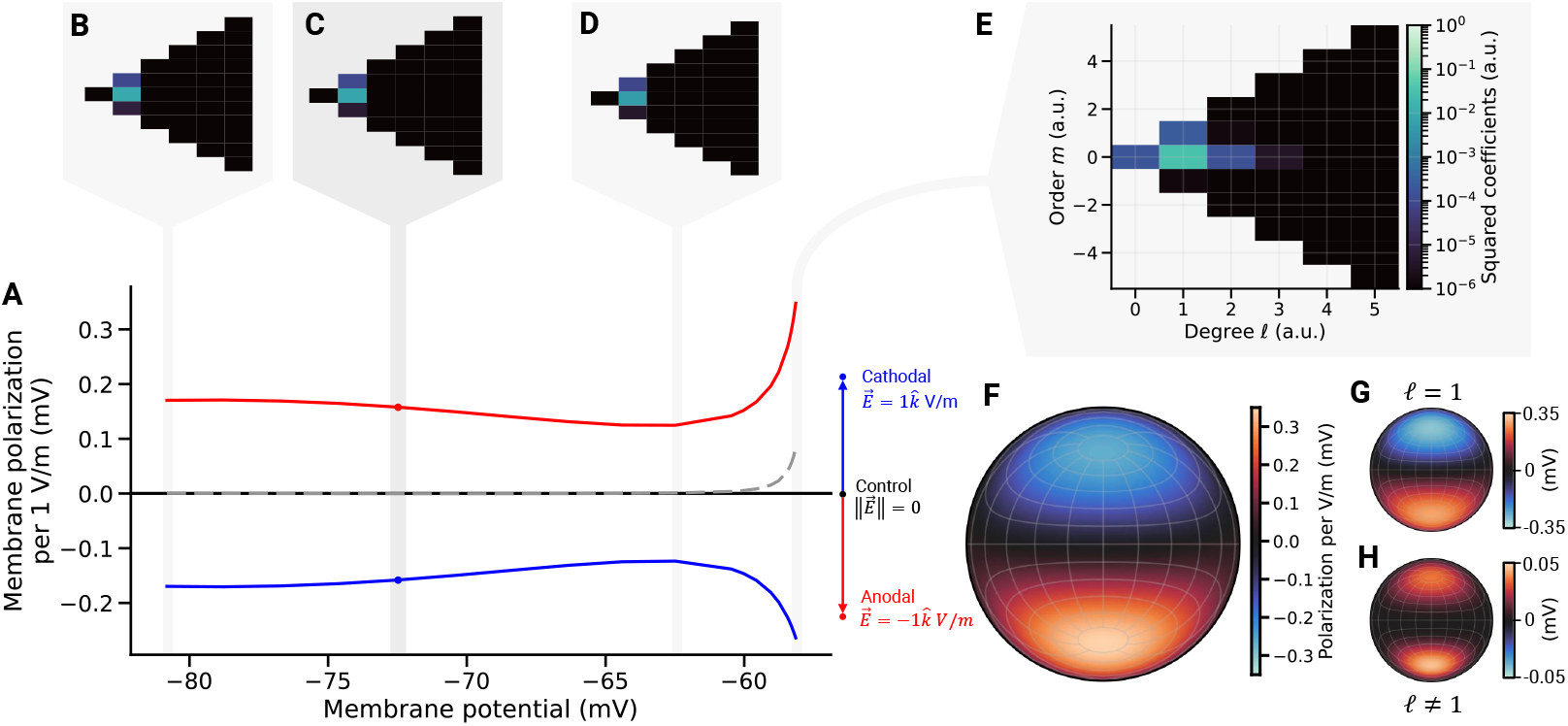
State dependence of somatic response to the electric field. **A.** Solid lines display the membrane polarization induced by an electric field of 1 V/m in anodal (red) and cathodal (blue) conditions. The control condition, i.e., zero-field, is displayed in black. The asymmetry between anodal and cathodal responses is displayed as the sum of both polarizations (dashed). The 2D spectrum of the response function is shown for hyperpolarization (**B**), resting (**C**), a depolarized state (**D**) and at the edge of the bifurcation (**E**). **F.** The response function at the edge of the bifurcation associated with E and its decomposition into dipolar (**G**, *ℓ* =1) and non-dipolar terms (**H**, *ℓ* ≠ 1).

### Reduced compartment models

The average response of a cell part composed of different compartments can be obtained by averaging the coefficients of all the compartments. In other words, averaging the coefficients provides a model of the average response of selected segments or cell types. This strategy can build an averaged response model of the axon, the apical dendrites, or other regions of interest depending on the experimental question. Regional averaged response models can be translated into multi-compartment models with just a few compartments representing the local membrane dynamics of a few parts of a cell, like the apical tuft, the soma, or the initial axon segment. Using averaged coefficients for each provides a statistic about the steady-state response to an external field in the specific region. Reducing the models here to just a few compartments allows us to use this knowledge in models at higher scales, like models of cortical columns made of multi-compartmental cells. Reducing cell models to just a few compartments significantly reduces the computation cost. The approach can be narrowed down to one-compartment models as long as they are assumed to be modeling a specific part of the cell. Then, it poses a proxy for stimulating commonly used models such as leaky or quadratic integrate and fire models [33].

### Implications for tDCS-based therapies

State-of-the-art montage optimization algorithms rely on the hypothesis that tDCS effects are related to the normal component of the electric field on the cortical surface. The present work cannot in itself be used as a direct input for optimization of electrode montages, as many other factors need to be considered. Nevertheless, it provides a direct translation between electric field and steady-state membrane potential change, see Appendix C. However, there is growing interest in using non-invasive brain stimulation techniques as a potential treatment for drug-resistant patients and using computational models to optimize these treatments [34]. An interesting role for computational models is to represent the pathophysiology underlying neurological disorders such as epileptic activity. For instance, in [35] and more recently in [36], neural mass models (NMM) have been used to represent epileptic seizure transitions: changes from interictal to ictal state were achieved by increasing the excitatory synaptic gain at the level of glutamatergic pyramidal cells and varying the inhibitory synaptic gains of GABAergic interneurons. Still, there is an open debate in the field about the validity of the predictions of such types of models [37]. Ideally, building pathophysiology models that can predict the effects on the pathology of induced electric fields in different brain regions is what will ultimately enable the definition of neurophysiologically relevant target functions. We believe that these will require models of the effects of electric fields that capture the behavior at the local population level by representing the different types of neurons and the specific effects of external fields on each. Our work represents a step in this direction.

### Translation to mesoscale models

The methods described are a starting point to derive the effects of weak, uniform electric fields at the population level (cortical column or small cortical patch scale). However, the translation from single cells to the mesoscale is not trivial, given that network effects are arising at the population level [15, 16]. A first approach can be to computationally obtain the response of a population by building models of cortical columns or networks of cell models and then extracting the effects at the population level. One could use a full cortical column model with multicompartment models such as the one developed by the Blue Brain Project [19] and include the effect of the field at each compartment of each cell. A less computationally expensive approach could be to build a similar cortical column model made of neuron models with reduced numbers of compartments, as introduced in the previous paragraph [38]. An alternative to computationally assessing the population-scale effects is to work in a theoretical framework that connects the microscale to the mesoscale, starting from simplifying assumptions. This kind of theoretical connection has recently grown in interest in the field of NMMs since the work by [39], who derived an exact mean field theory from the quadratic integrate and fire neuron model. In this framework, values for the induced transmembrane currents by weak fields are especially relevant since there is no ambiguity in translating the effects from the microscale to the population level.

## 5. Conclusions

In this work, we have provided a framework for analysis of the effects of weak, uniform electric fields in neuron models in terms of changes in their steady-state conditions. We quantify the induced perturbation dependence on the orientation using spherical harmonic coefficients. Through this analysis, we have validated numerically the commonly used dipole approximation for the studied models in their resting state (far from the threshold) and weak electric field regime. We have also provided a theoretical justification for this in a generic passive membrane model. We have shown, however, that there are conditions under which this approximation is less accurate—particularly close to the spiking threshold. We have also seen that although pyramidal cells generally display the most substantial effects, some interneurons can also be strongly affected but with a more diverse set of responses. Although the impact of these effects on neuron networks is left for future research, our work provides the basis for further computational studies studying the relation of electric field effects at microscale and mesoscales. This can benefit the development of brain models and derived clinical applications relying on model-driven brain stimulation with weak electric fields, such as tES.

## Acknowledgments

This work has received funding from the European Research Council (ERC) under the European Union’s Horizon 2020 research and innovation programme (grant agreement No 855109; ERC-SyG 2019 Galvani) and from FET under the European Union’s Horizon 2020 research and innovation programme (grant agreement No 101017716, Neurotwin).

## Appendix A. Real spherical harmonics

A square-integrable function 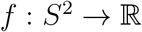 can be expanded in terms of the real harmonics 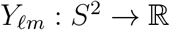 like:

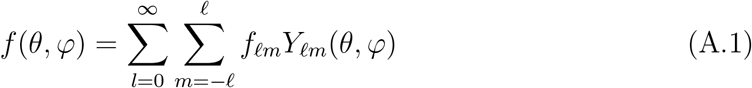

The general definition of each spherical harmonic function *Y_ℓm_* can be cumbersome, and its derivation is out of the scope of this text. We summarize how they can be stated for completeness, but it does not serve as an introduction to the field. Here it is worthwhile to recall the notation:

- *θ* stands for the polar angle or inclination.
- *φ* for the azimuth or longitude.
- *ℓ* for the spherical harmonic degree.
- *m* for the order.

Under this convention, the real spherical harmonics are defined as [24]:

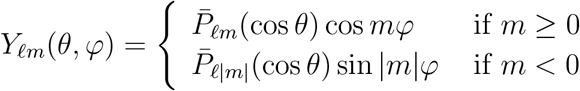

where 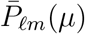 are the normalized associated Legendre functions. For 4*π*-normalized spherical harmonic functions, they can be written as:

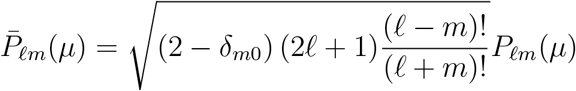

Note that *δ_ij_* refers to the Kronecker delta and *P_ℓm_*(*μ*) refers to the unnormalized associated Legendre functions. They can be derived from the standard Legendre polynomials using the relations shown below:

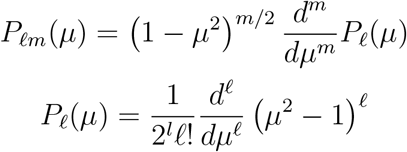

As a set of basis functions, they need to be orthogonal. Note that the associated Legendre functions are indeed orthogonal for any given m, given that:

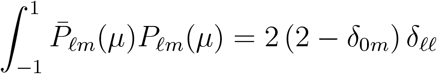

## Appendix B. Dipole nature of electric field effect in compartment model of a neuron in the passive, linear membrane regime

Here we show that for a passive membrane, the effect of an electric field on a neuron compartment, or its average over a set of compartments (uniform field), can be written in the form *δV* = **P** · **E**, where **P** is a vector. Note that the behavior far from the threshold and under weak electric fields is considered comparable to that of a passive membrane since the active properties of membranes (nonlinear) require more significant effects to come into play. This is the so-called “λE” model used in transcranial electrical stimulation models [5, 40].

The effect of an external field can be represented in a neuron compartment model by an axial current that results from the potential difference induced by the field along the fiber associated with a compartment

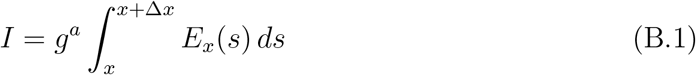

where *g^a^* is the axial conductivity and *E_x_* is the field along the fiber. We can rewrite this as *I* = *g^a^ ∫_f_* **E** · *d*l with the line integral along the fiber *f* compartment. If the field is constant along the fiber we can express this simply as *I* = *g^a^* **E** · **u**. The vector **u** is a vector defined from one center of compartment to another center of a connected compartment, and pointing into the compartment of interest (see Figure B1). If the E field is aligned with this direction, we get a positive current into the compartment. If the compartment of interest is *k* and the connected compartment is *j*, we refer to the vector **u***_k,j_* parallel to **x***_k_* – **x***_j_* (the coordinates of the compartments).

**Figure B1.**
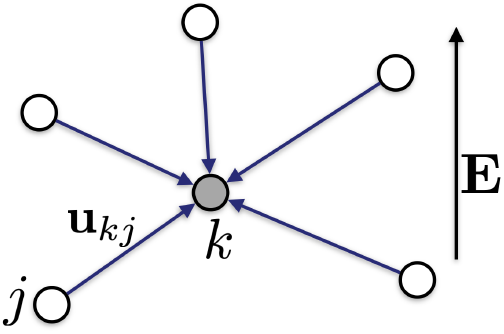
Compartments and vectors for cable equation in E field. Here we represent a fiber by a node and a set of directed edges.

From conservation of charge (the sum over j is over connected compartments, and *i* over membrane currents), and assuming that the field is uniform and thus independent of the compartment *k*, we obtain

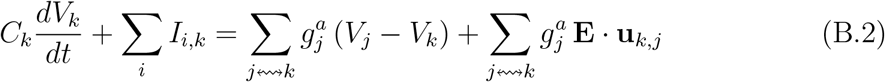

where the sum on the left hand side is over ionic membrane currents out of the compartment. The notation 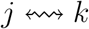 indicates a sum over all compartments *j* connected to the *k*th one. For an arbitrary compartment-dependent quantity *Q_j_*, sums over 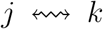 can be rewritten via a compartment connectivity matrix *c_kj_*, with value of 1 if compartment *j* is connected to *k*, zero otherwise. That is, 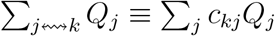.

We can simplify this and also pull the constant electric field out the sum,

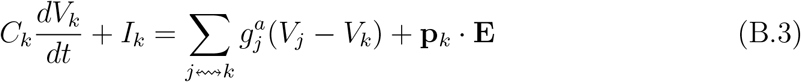

where *I_k_* = ∑*_i_ I_i,k_* and where

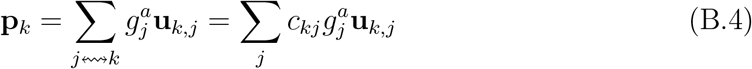

The equation for a single compartment in steady state is thus of the form

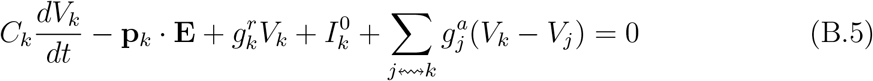

where **p***_k_* is the dipole vector corresponding to the *k*th compartment, and 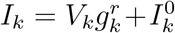, with 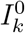 the sum of currents associated to the reversal potentials, and 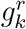 the radial (transmembrane) conductivity in the linear regime.

Thus, we see that the behavior of each compartment is characterized by its own dipole and the interaction with other compartments.

Equation B.5 is linear and can be expressed in matrix form, with the compartment array notation **V** = (*V_k_*) = (*V*_1_, *V*_2_,…, *V_N_*)*^T^, C* = diag(*C_k_*) = diag(*C*_1_, *C*_2_,…, *C_N_*), and *P* = (**p**_1_,…, **p***_N_*)*^T^*

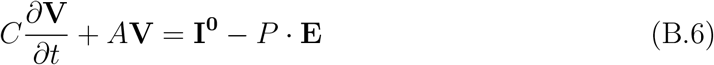

and with the matrix *A* given by

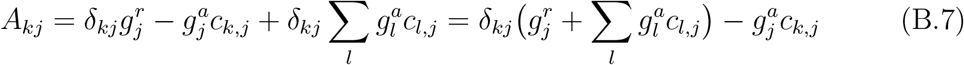

where we split the diagonal and off diagonal elements in the second equality.

We recall here that a matrix is strictly diagonally dominant if, for every row of the matrix, the magnitude of the diagonal entry in a row is larger than or equal to the sum of the magnitudes of all the other (non-diagonal) entries in that row, i.e.,

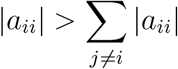

Now, the Gershgorin circle theorem implies that a strictly diagonally dominant matrix (or an irreducibly diagonally dominant matrix) is non-singular [41] [Th 6.2.27]. We can observe that this is the case for A, since (all g’s are taken to be positive, and recall the *c*’s are 1s or 0s, with *c_ii_* = 0)

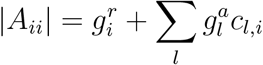

and

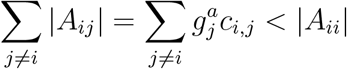

This means that the matrix *A* has an inverse.

In steady state 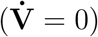, the solution to this equation is

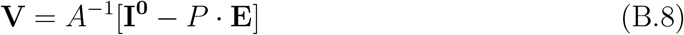

or

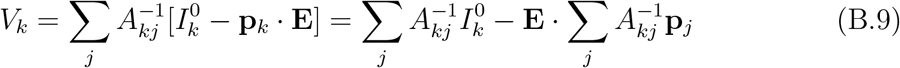

where the second equality is a result of linearity of the *A* matrix, i.e., the fact that we are dealing with a linear equation. Finally, we can write the solution as

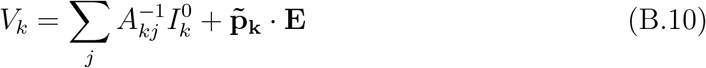

This equation shows that the response of the compartment to an electric field is of dipole form, with the total dipole 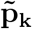 resulting from a superposition of dipoles,

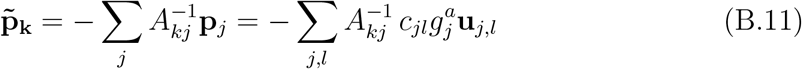

Furthermore, we observe that any linear function of compartment potentials will also display, by further superposition, a dipole response. This includes, for example, the average membrane perturbation for apical dendrite compartments, or soma.

What happens in the **nonlinear case**, when the neuron is not at baseline? We can write (steady state)

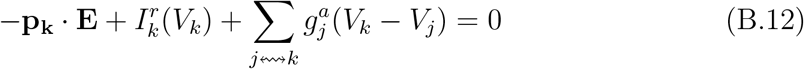

with 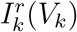 a nonlinear function, or, in a fashion analogous to the discussion above, in matrix form

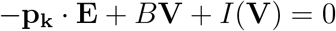

or

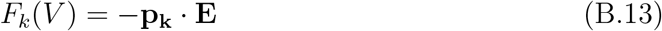

for a nonlinear operator *F*. This means we cannot pull out the E field out of the inverse. However, if the electric field is very weak, we can carry out the analysis above with the linearized version of Equation B.13 by expanding it around the potential with zero field, with the same conclusion.

## Appendix C. Realistic montage exploration

### Context

Based on the *λ*E approach, one logical strategy is to optimize for the normal component of the electric field with respect to the cortical surface. The quantification of the performance of the montage is carried out by averaging the electric field on the target area. Expressions like the *λ*E model, which captures the main response of neurons to the fields can be used to develop metrics that are more representative of the relevant physiological effects of the stimulation. As an example of this, we performed a montage optimization to stimulate the dorsolateral prefrontal cortex (dlPFC), which is a common area for stimulation in tDCS, in a real head model - see Figure C1.A.

### Montage creation

We ran the Stimweaver algorithm [40] with a maximum of 8 channels, and a total injected current of 4.0 mA, under the restriction of 2.0 mA maximum per channel. The montage was optimized to achieve a normal component to the cortical surface of the electric field of 0.25 V/m in the dlPFC, using weights of 10 and 2 to on- and off-target nodes, respectively. We computed the predicted average membrane polarization of LV pyramidal cells somata on- and off-target as a proof of concept (Figure C1).

**Figure C1.**
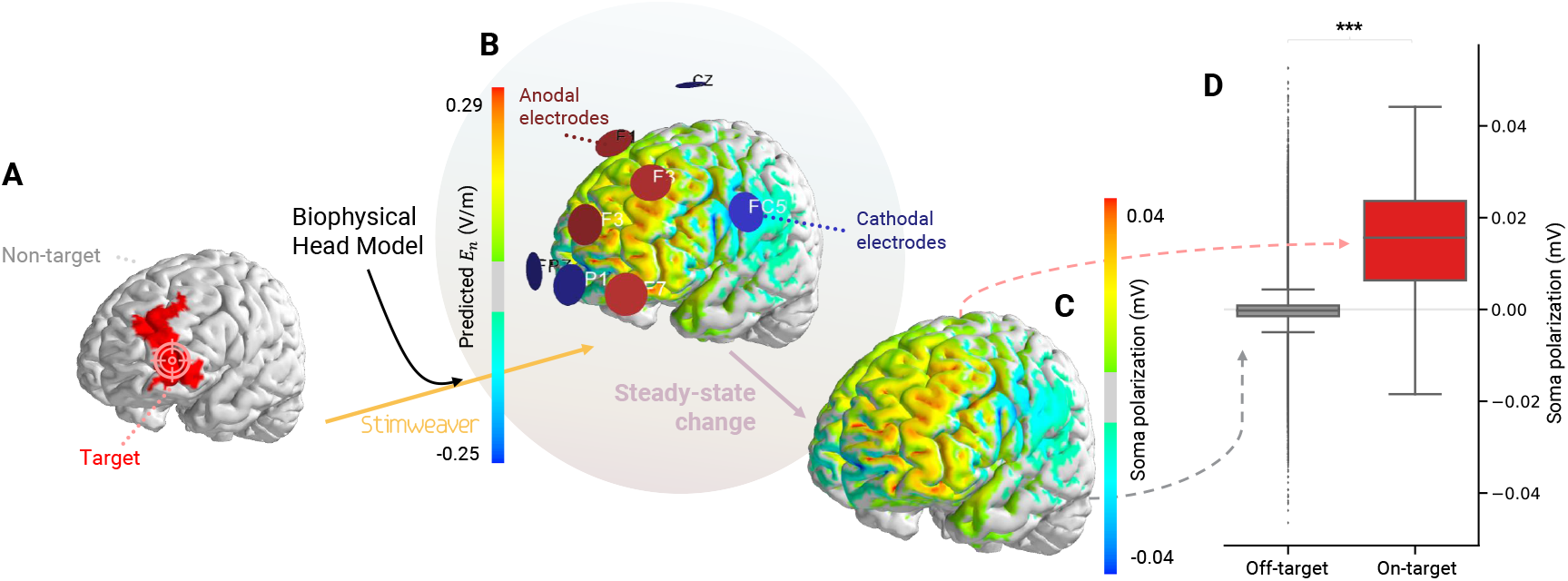
Predicted effects in a realistic montage. Representation of a pipeline from a target definition to a predicted physiological effect. **A: Target definition.** Overlay in red of the cortical area marked as the target of the stimulation. In gray, the rest of the cortex. **B: Optimized montage.** Anodal (blue) and cathodal (red) electrodes location over the cortical surface. The predicted electric field magnitude is displayed color coded over the cortical surface, being the color code the one displayed in the colorbar on the left (V/m). **C: Predicted polarization.** Prediction of the somatic polarization of LV thick-tufted pyramidal cells. **D: Quantification of effects on and off target.** Predicted values of steady-state somatic polarization induced in a LV thick-tufted pyramidal cell using the optimized montage shown in B. The figure shows how effects are induced in the cortex selectively to the original target.

### Results

The target map of the montage is shown in Figure C1.A. The optimized montage is shown in Figure C1.B, together with the predicted electric fields on the cortex—overlaid on the brain morphology. The normal component on-target predicted with the optimized montage has a median value across nodes of 0.11 V/m and a mode of 0.17 V/m computed as the maximum of the kernel density estimate. Through the formalism in Equation 2 and the computationally derived response functions, that map can be translated to a variable with actual physiological meaning, e.g., the somatic membrane potential perturbation of LV early branching thick-tufted pyramidal cells, Figure C1.C. We observe a significant effect on-target (Mann-Whitney U, p< 1 × 10^-3^, see Figure C1.D).

## Footnotes

§ More generally, we can start from 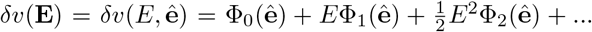 The first term is zero here by definition.

## References

[1] M. A. Nitsche and W. Paulus. Excitability changes induced in the human motor cortex by weak transcranial direct current stimulation. J Physiol, 527 Pt 3:633–9, 2000.

[2] Hebb, d. o. the organization of behavior: A neuropsychological theory. Science Education, 34(5):336–337, December 1949.

[3] D. Liebetanz, F. Fregni, K. K. Monte-Silva, M. B. Oliveira, A. Amancio-dos Santos, M. A. Nitsche, and R. C. Guedes. After-effects of transcranial direct current stimulation (tDCS) on cortical spreading depression. Neurosci Lett, 398(1-2):85–90, 2006.

[4] D. Liebetanz, M. A. Nitsche, F. Tergau, and W. Paulus. Pharmacological approach to the mechanisms of transcranial DC-stimulation-induced after-effects of human motor cortex excitability. Brain, 125(2238-2247), 2002.

[5] Giulio Ruffini, Fabrice Wendling, Isabelle Merlet, Behnam Molaee-Ardekani, Abeye Mekkonen, Ricardo Salvador, Aureli Soria-Frisch, Carles Grau, Stephen Dunne, and Pedro Miranda. Transcranial Current Brain Stimulation (tCS): Models and Technologies. IEEE Transactions on Neural Systems and Rehabilitation Engineering, 21(3):333–345, May 2013.

[6] Pedro Cavaleiro Miranda, Abeye Mekonnen, Ricardo Salvador, and Giulio Ruffini. The electric field in the cortex during transcranial current stimulation. Neuroimage, 70:45–58, 2013.

[7] P. C. Miranda, M. Hallett, and P. J. Basser. The electric field induced in the brain by magnetic stimulation: a 3-D finite-element analysis of the effect of tissue heterogeneity and anisotropy. IEEE Trans Biomed Eng, 50(9):1074–85, 2003.

[8] Frank Rattay. Analysis of models for external stimulation of axons. IEEE Transactions on Biomedical Engineering, 33(10):974–977, 1986.

[9] Thomas Radman, Raddy L. Ramos, Joshua C. Brumberg, and Marom Bikson. Role of cortical cell type and morphology in subthreshold and suprathreshold uniform electric field stimulation in vitro. Brain Stimulation, 2(215):28, 2009.

[10] W. Rall. Core conductor theory and cable properties of neurons. pages 39–97. American Physiological Society, 1977.

[11] D. Tranchina and C. Nicholson. A model for the polarization of neurons by extrinsically applied electric fields. Biophys J, 50(6):1139–56, 1986.

[12] Frank Rattay. Analysis of models for extracellular fiber stimulation. IEEE Transactions on Biomedical Engineering, 36(7):676–682, 1989.

[13] B. J. Roth. Mechanisms for electrical stimulation of excitable tissue. Crit Rev Biomed Eng, 22(3-4):253–305, 1994.

[14] Aman S Aberra, Angel V Peterchev, and Warren M Grill. Biophysically realistic neuron models for simulation of cortical stimulation. Journal of Neural Engineering, 15(6):066023, October 2018.

[15] F Fröhlich and D. A. McCormick. Endogenous Electric Fields May Guide Neocortical Network Activity. Neuron, 67:129–143, 2010.

[16] D. Reato, A. Rahman, M. Bikson, and L. C. Parra. Low-intensity electrical stimulation affects network dynamics by modulating population rate and spike timing. Journal of Neuroscience, 30(45):15067–15079, November 2010.

[17] Pierre Simon Laplace, Nathaniel Bowditch, and N. I Bowditch. Mécanique céleste. Hillard, Gray, Little, and Wilkins, 1829.

[18] W.T.B. Kelvin and P.G. Tait. Treatise on Natural Philosophy. Number v. 1 in Clarendon Press series. Clarendon Press, 1867.

[19] Henry Markram, Eilif Muller, Srikanth Ramaswamy, Michael W. Reimann, Marwan Abdellah, Carlos Aguado Sanchez, Anastasia Ailamaki, Lidia Alonso-Nanclares, Nicolas Antille, Selim Arsever, Guy Antoine Atenekeng Kahou, Thomas K. Berger, Ahmet Bilgili, Nenad Buncic, Athanassia Chalimourda, Giuseppe Chindemi, Jean-Denis Courcol, Fabien Delalondre, Vincent Delattre, Shaul Druckmann, Raphael Dumusc, James Dynes, Stefan Eilemann, Eyal Gal, Michael Emiel Gevaert, Jean-Pierre Ghobril, Albert Gidon, Joe W. Graham, Anirudh Gupta, Valentin Haenel, Etay Hay, Thomas Heinis, Juan B. Hernando, Michael Hines, Lida Kanari, Daniel Keller, John Kenyon, Georges Khazen, Yihwa Kim, James G. King, Zoltan Kisvarday, Pramod Kumbhar, Sébastien Lasserre, Jean-Vincent Le Bé, Bruno R.C. Magãlhaes, Angel Merchán-Pérez, Julie Meystre, Benjamin Roy Morrice, Jeffrey Muller, Alberto Muñoz-Céspedes, Shruti Muralidhar, Keerthan Muthurasa, Daniel Nachbaur, Taylor H. Newton, Max Nolte, Aleksandr Ovcharenko, Juan Palacios, Luis Pastor, Rodrigo Perin, Rajnish Ranjan, Imad Riachi, José-Rodrigo Rodríguez, Juan Luis Riquelme, Christian Rössert, Konstantinos Sfyrakis, Ying Shi, Julian C. Shillcock, Gilad Silberberg, Ricardo Silva, Farhan Tauheed, Martin Telefont, Maria Toledo-Rodriguez, Thomas Tränkler, Werner Van Geit, Jafet Villafranca Díaz, Richard Walker, Yun Wang, Stefano M. Zaninetta, Javier DeFelipe, Sean L. Hill, Idan Segev, and Felix Schürmann. Reconstruction and simulation of neocortical microcircuitry. Cell, 163(2):456–492, October 2015.

[20] Cameron C. McIntyre and Warren M. Grill. Extracellular stimulation of central neurons: Influence of stimulus waveform and frequency on neuronal output. Journal of Neurophysiology, 88(4):1592–1604, October 2002.

[21] M. L. Hines and N. T. Carnevale. NEURON: a tool for neuroscientists. Neuroscientist, 7(2):123–35, 2001.

[22] Huan Wang, Bonnie Wang, Kieran P. Normoyle, Kevin Jackson, Kevin Spitler, Matthew F. Sharrock, Claire M. Miller, Catherine Best, Daniel Llano, and Rose Du. Brain temperature and its fundamental properties: a review for clinical neuroscientists. Frontiers in Neuroscience, 8, October 2014.

[23] J.R. Driscoll and D.M. Healy. Computing fourier transforms and convolutions on the 2-sphere. Advances in Applied Mathematics, 15(2):202–250, June 1994.

[24] Mark A. Wieczorek and Matthias Meschede. SHTools: Tools for working with spherical harmonics. Geochemistry, Geophysics, Geosystems, 19(8):2574–2592, August 2018.

[25] J. A. Rod Blais. Discrete spherical harmonic transforms: Numerical preconditioning and optimization. In Computational Science – ICCS 2008, pages 638–645. Springer Berlin Heidelberg, 2008.

[26] M. Bikson, M. Inoue, H. Akiyama, J. K. Deans, J. E. Fox, H. Miyakawa, and J. G. Jefferys. Effects of uniform extracellular DC electric fields on excitability in rat hippocampal slices in vitro. J Physiol, 557(Pt 1):175–90, 2004.

[27] Beatriz Rebollo, Bartosz Telenczuk, Alvaro Navarro-Guzman, Alain Destexhe, and Maria V. Sanchez-Vives. Modulation of intercolumnar synchronization by endogenous electric fields in cerebral cortex. Science Advances, 7(10), March 2021.

[28] Walter Paulus and John C. Rothwell. Membrane resistance and shunting inhibition: where biophysics meets state-dependent human neurophysiology. The Journal of Physiology, 594(10):2719–2728, May 2016.

[29] Marom Bikson and Asif Rahman. Origins of specificity during tDCS: anatomical, activity-selective, and input-bias mechanisms. Frontiers in Human Neuroscience, 7, 2013.

[30] T. Radman, Y. Su, J. H. An, L. C. Parra, and M. Bikson. Spike timing amplifies the effect of electric fields on neurons: implications for endogenous field effects. J Neurosci, 27(11):3030–6, 2007.

[31] Daniel E. Feldman. The spike-timing dependence of plasticity. Neuron, 75(4):556–571, August 2012.

[32] Greg Kronberg, Asif Rahman, Mahima Sharma, Marom Bikson, and Lucas C. Parra. Direct current stimulation boosts hebbian plasticity in vitro. Brain Stimulation, 13(2):287–301, March 2020.

[33] Peter Dayan and L F Abbott. Theoretical neuroscience. Computational neuroscience. MIT Press, London, England, December 2001.

[34] Giulio Ruffini, Fabrice Wendling, Roser Sanchez-Todo, and Emiliano Santarnecchi. Targeting brain networks with multichannel transcranial current stimulation (tCS). Current Opinion in Biomedical Engineering, 2018.

[35] F. Wendling, F. Bartolomei, J. J. Bellanger, and P. Chauvel. Epileptic fast activity can be explained by a model of impaired GABAergic dendritic inhibition. Eur J Neurosci, 15(9):1499–508, 2002.

[36] Edmundo Lopez-Sola, Roser Sanchez-Todo, Elia Lleal, Elif Köksal Ersöz, Maxime Yochum, Julia Makhalova, Borja Mercadal, Maria Guasch, Ricardo Salvador, Diego Lozano-Soldevilla, Julien Modolo, Fabrice Bartolomei, Fabrice Wendling, Pascal Benquet, and Giulio Ruffini. A personalizable autonomous neural mass model of epileptic seizures. Journal of Neural Engineering, 2022.

[37] Nicolás Deschle, Juan Ignacio Gossn, Prejaas Tewarie, Björn Schelter, and Andreas Daffertshofer. On the validity of neural mass models. Frontiers in Computational Neuroscience, 14, January 2021.

[38] Yves Denoyer, Isabelle Merlet, Fabrice Wendling, and Pascal Benquet. Modelling acute and lasting effects of tDCS on epileptic activity. Journal of Computational Neuroscience, 48(2):161–176, May 2020.

[39] Ernest Montbrió, Diego Pazó, and Alex Roxin. Macroscopic description for networks of spiking neurons. Phys. Rev. X, 021028, 2015.

[40] Giulio Ruffini, Michael D. Fox, Oscar Ripolles, Pedro Cavaleiro Miranda, and Alvaro Pascual-Leone. Optimization of multifocal transcranial current stimulation for weighted cortical pattern targeting from realistic modeling of electric fields. NeuroImage, 89:216–225, 2014.

[41] Roger A Horn and Charles R Johnson. Matrix Analysis. Cambridge University Press, Cambridge, England, February 1990.

